## Supplementary material for "Diversity in the internal functional feeding elements of sympatric morphs of Arctic charr (Salvelinus alpinus)": All supplemental materials

Affiliations

Corresponding authors:

*S1 Appendix: Sampling scheme and summary statistics for the four sympatric charr morphs (numbers of individuals, average fork length (FL), weight and age and the standard deviation (SD) for length, weight and age of the 240 individuals).*

| Morph | Sex | N | $FL_{Mean}$ (cm) | $Weight_{Mean}$ (g) | $Age_{Mean}$ |
|---|---|---|---|---|---|
| LB | F | 17 | 42.28 ± 7.01 | 944.29 ± 381.73 | 8.20 ± 1.61 |
| | M | 32 | 40.73 ± 6.33 | 844.66 ± 388.13 | 8.06 ± 2.20 |
| | All | 49 | 41.27 ± 6.54 | 879.23 ± 384.91 | 8.11 ± 2.01 |
| SB | F | 47 | 16.46 ± 4.36 | 59.77 ± 56.00 | 5.60 ± 1.36 |
| | M | 33 | 13.79 ± 2.88 | 30.91 ± 23.05 | 5.34 ± 2.04 |
| | Unknown | 2 | 14.70 ± 1.70 | 34.36 ± 13.38 | 4.00 ± 0.00 |
| | All | 82 | 15.34 ± 3.97 | 47.54 ± 46.87 | 5.46 ± 1.66 |
| PL | F | 23 | 22.47 ± 3.30 | 106.49 ± 59.02 | 6.87 ± 1.22 |
| | M | 38 | 16.75 ± 3.31 | 52.90 ± 33.01 | 5.14 ± 1.00 |
| | All | 61 | 18.91 ± 4.31 | 73.10 ± 51.33 | 5.80 ± 1.38 |
| PI | F | 14 | 38.25 ± 6.05 | 670.43 ± 337.38 | 8.93 ± 1.73 |
| | M | 34 | 38.19 ± 8.56 | 780.21 ± 597.28 | 8.61 ± 1.52 |
| | All | 48 | 38.20 ± 7.84 | 748.19 ± 533.39 | 8.70 ± 1.57 |
| All | F | 101 | 25.19 ± 11.61 | 303.94 ± 407.21 | 6.78 ± 1.90 |
| | M | 137 | 26.96 ± 13.38 | 413.04 ± 520.13 | 6.74 ± 2.32 |
| | All | 240 | 26.11 ± 12.66 | 363.97 ± 476.59 | 6.73 ± 2.16 |

[1]Fork length: length from tip of snout to the posterior tip of middle caudal fin ray

Log: natural logarithmic transformation.

*S2 Appendix: Explanation of 37 landmarks used for landmarking the external shape, X-S indicates sliding (semi) landmark.*

| Landmark(s) | Details on placement and rationale |
|---|---|
| 1, 2, 3, 4 and 22 | Capture the curvature of the snout: 1 – intersection of upper jaw and snout; 2 – anterior-most point of snout; 3 – inflection of snout; 4 – notch of frontal bone; 22 – anterior-most point of lower jaw |
| 5 | Posterior end of skull, most postero-dorsal point of supraoccipital |
| 6-S | Middle between LM 5 and 7, on predorsal surface |
| 7 and 8 | Location of insertion of the Dorsal fin (7 - anterior and 8- posterior) |
| 9 and 10 | Location of insertion of the Adipose fin (9 - anterior and 10- posterior) |
| 11 and 14 | Characterizes caudal peduncle depth. 11 – inflection of curve between adipose and Caudal fin (dorsal); 14 – inflection of curve between Anal fin and Caudal fin (ventral) |
| 12 | Posterior-most tip of the middle Caudal fin rays. Used to measure the fork length (FL) of the fish when paired with LM 2 |
| 13 | End of Lateral line. Used to measure the standardized length (SL) of the fish, when paired with LM 2 |
| 15 and 16 | Location of insertion of the Anal fin (15 – posterior and 16 – anterior) |
| 17-S | Location of anterior insertion of the Pelvic fin |
| 18-S and 19-S | Characterizes the curve of the ventral surface between the head and Pelvic fin. 18 – point of max body depth. 19 – midpoint between 18 and 20 |
| 20 and 21 | Location of the branchiostegal rays (20 – posterior and 21 – anterior) |
| 23 | Location of center of pupil |
| 24-S, 25-S, 26-S and 27-S | Characterize the eye location and size, landmarks placed on the orbital bones (24 – dorsal margin; 25 - posterior margin; 26 – ventral margin; 27 – anterior margin) |
| 1, 28, 29 and 30 | Characterizes the Maxilla.1 -anterior-most point of the bone; 28 - dorsal surface of the bone just beneath the eye; 29 - perpendicular to 28 on the ventral surface; 30 – posterior-most point of the bone |
| 31 | Location of the Preopercular bone, dorsal-most point of the bone |
| 32 and 33 | Location of the Opercular (dorsal-most point) and Subopercular bones (dorsal-most point) |
| 34, 35 and 36 | Characterizes the Lateral line, placed in reference to points on the back (34 is below 7, 35 below 8 and 36 below 9). Perpendicular to midline |
| 37 | Location of the Pectoral fin, dorso-anterior insertion |

*S3 Appendix: Explanation of landmarks used to capture the shape of the six bones (dentary, articular-angular, quadrat, premaxilla, maxilla and supramaxilla). X-S indicates sliding landmark. See Fig 1D - E for placement of landmarks on bones.*

| Bones | Landmark(s) | Details on placement and rationale |
|---|---|---|
| Dentary | 1 | Ventral tip of the mandibular symphysis. |
| | 2 | Anterior lingual palate, at the anterior base of the anterior-most tooth. |
| | 3 | Middle between LM 2 and 4, along surface of dorsal surface of lingual palate |
| | 4 | Posterior lingual palate, at the posterior base of the most posterior tooth |
| | 5 | Anterior base of the coronoid process. Along with LM 6, 7 and 8, used to characterize the coronoid process |
| | 6 | Dorso-anterior tip of coronoid process. Along with LM 5, 7 and 8, used to characterize the coronoid process |
| | 7 | Dorso-posterior tip of coronoid process. Along with LM 5, 6 and 8, used to characterize the coronoid process. |
| | 8 | Ventro-posterior tip of coronoid process. Along with LM 5, 6 and 7, used to characterize the coronoid process |
| | 9 | Inflection of the posterior margin of the medial wall, between LM 8 and 11 |
| | 10-S | Intersection of medial wall and ventral ridge, location on inflection between LM 9 and 11 |
| | 11,12 and 13 | Used to characterize the shape of the most Posterior part of the ventral ridge and shelf, 11 - dorso-posterior tip of ventral shelf; 12 – posterior-most tip of ventral shelf; 13 - ventro-posterior tip of ventral shelf |
| | 14-S | Capturing the ventral curve of the ventral shelf, placed in middle between LM 1 and 13 |
| | 15-S | Inflection of ventral shelf between LM 1 and 14 |
| | 16 | Anterior intersection of the dorsal and ventral ridges |
| Articular - angular | 1 | Anterior-most tip of anterior process |
| | 2 | Inflection of curvature of dorsal margin of anterior process toward coronoid process. Characterize curvature of the anterior process dorsal margin, together with LM 3 |
| | 3 | Intersection of anterior process dorsal margin and coronoid process. Characterizes curvature of the anterior process dorsal margin, together with LM 2 |
| | 4 | Anterior-most tip of coronoid process |
| | 5 | Base of coronoid process, where curve changes between LM 4 and 6 |
| | 6 | Anterior dorsal tip of quadrate facet |
| | 7 | Ventral-most point in depression of quadrate facet |
| | 8 | Posterior dorsal tip of quadrate facet |

| | | |
|---|---|---|
| | 9 | Posterior-most point of retroarticular |
| | 10 | Ventro-posterior tip of retroarticular |
| | 11 | Anterior process of retroarticular |
| | 12 | Intersection between anterior process of retroarticular and anterior process of Articular-angular |
| | 13-S | Minor inflection on ventral surface of anterior process. Characterizes ventral surface curvature of anterior process, together with LM 14. |
| | 14-S | Main inflection on ventral surface of anterior process. Characterizes ventral surface curvature of anterior process, together with LM 13. |
| | 15 | Anterior tip of the quadrate facet, at the base of the Articular-angular 'wing' (thin ossification connecting anterior process and coronoid process) |
| Quadrat | 1 | Anterior-most tip of the dorsal margin |
| | 2 | Middle between LM 1 and 3 |
| | 3 | Posterior-most tip of the dorsal margin |
| | 4-S | Inflection of the symplectic incisure (intersection of body and preopercular process) |
| | 5 | Posterior-most tip of preopercular process |
| | 6 | Ventral base of the preopercular process |
| | 7 | Inflection of the preopercular base, ventro-posterior-most point between LM 6 and 8 |
| | 8 | Intersection of lateral condyle and base of preopercular process, |
| | 9 | Ventral-most point on lateral condyle |
| | 10 | Intersection of lateral and medial condyles |
| | 11 | Anterior-most tip of medial condyle |
| | 12 | Intersection of body and medial condyle |
| | 13 | Intersection of body and lateral condyle, middle point between LM 10 and 12, placed on the body. |
| Premaxilla | 1 | Ventral surface of the posterior limb, placed at the posterior base of the posterior-most tooth |
| | 2 | Dorso-posterior tip of the posterior limb |
| | 3 | Ventral inflection at the intersection of the posterior limb and ascending limb |
| | 4 | Dorsal inflection at the intersection of the posterior limb and ascending limb |
| | 5 | Posterior tip of the dorsal margin of the ascending limb |
| | 6 | Dorsal-most tip of the ascending process |
| | 7 | Anterior tip of the dorsal margin of the ascending limb |
| | 8 | Intersection of the ascending limb and anterior limb |
| | 9 | Dorsal tip of premaxillary symphysis |

| | | |
|---|---|---|
| | 10 | Ventral surface of the anterior limb, placed at the anterior base of the anterior-most tooth |
| | 11 | Ventral surface of the posterior limb, perpendicular to LM 6 |
| | 12 | Ventral surface of anterior limb, middle between LM 1 and 10 |
| Maxilla | 1 | Dorsal inflection of the caudal lobe, between LM 2 and 5 Characterizes the curve of the caudal lobe, together with LM 2 and 3. |
| | 2 | Posterior-most tip of the caudal lobe. Characterizes the curve of the caudal lobe, together with LM 1 and 3. |
| | 3 | Ventral inflection of the caudal lobe, between LM 2 and 4. Characterizes the curve of the caudal lobe, together with LM 1 and 2. |
| | 4 | Posterior dental palate, at the posterior base of the most posterior tooth |
| | 5 | Point perpendicular to LM 4 on the dorsal surface of maxillary body |
| | 6 | Middle between LM 4 and 8, along dental palate |
| | 7 | Point perpendicular to LM 6 on the dorsal surface of maxillary body |
| | 8 | Anterior dental palate, at anterior base of the most anterior tooth |
| | 9 | Ventral base of maxillary neck. Characterizes the curves of maxillary head and neck, together with LM 10,11,12,13 and 14. |
| | 10 | Ventral inflection of the maxillary neck. Characterizes the curves of the maxillary head and neck, together with LM 9,11,12,13 and 14. |
| | 11 | Anterior-most point of the maxillary head. Characterizes the curves of the maxillary head and neck, together with LM 9,10,12,13 and 14 |
| | 12 | Dorsal inflection of the maxillary head. Characterizes the curves of the maxillary head and neck, together with LM 9,10,11,13 and 14. |
| | 13 | Dorsal inflection of the maxillary neck. Characterizes the curves of the maxillary head and neck, together with LM 9,10,11,12 and 14. |
| | 14 | Dorsal base of the maxillary neck. Characterizes the curves of the maxillary head and neck, together with LM 9,10,11,12 and 13. |
| | 15-S and 16-S | Characterize curvature of the dorsal ridge between LM 14 and 7. Both landmarks placed on curve inflections |
| | 17-S | Inflection of the dorsal ridge between LM 7 and 5 |
| Supramaxilla | 1 | Anterior-most tip of the bone |

| | | |
|---|---|---|
| | 2 | Anterior base of the ventral protrusion |
| | 3 | Ventral-most tip of the ventral protrusion |
| | 4 | Posterior base of the ventral protrusion |
| | 5 | Posterior-most tip of the bone |
| | 6 | Point perpendicular to LM 3 on the dorsal surface |

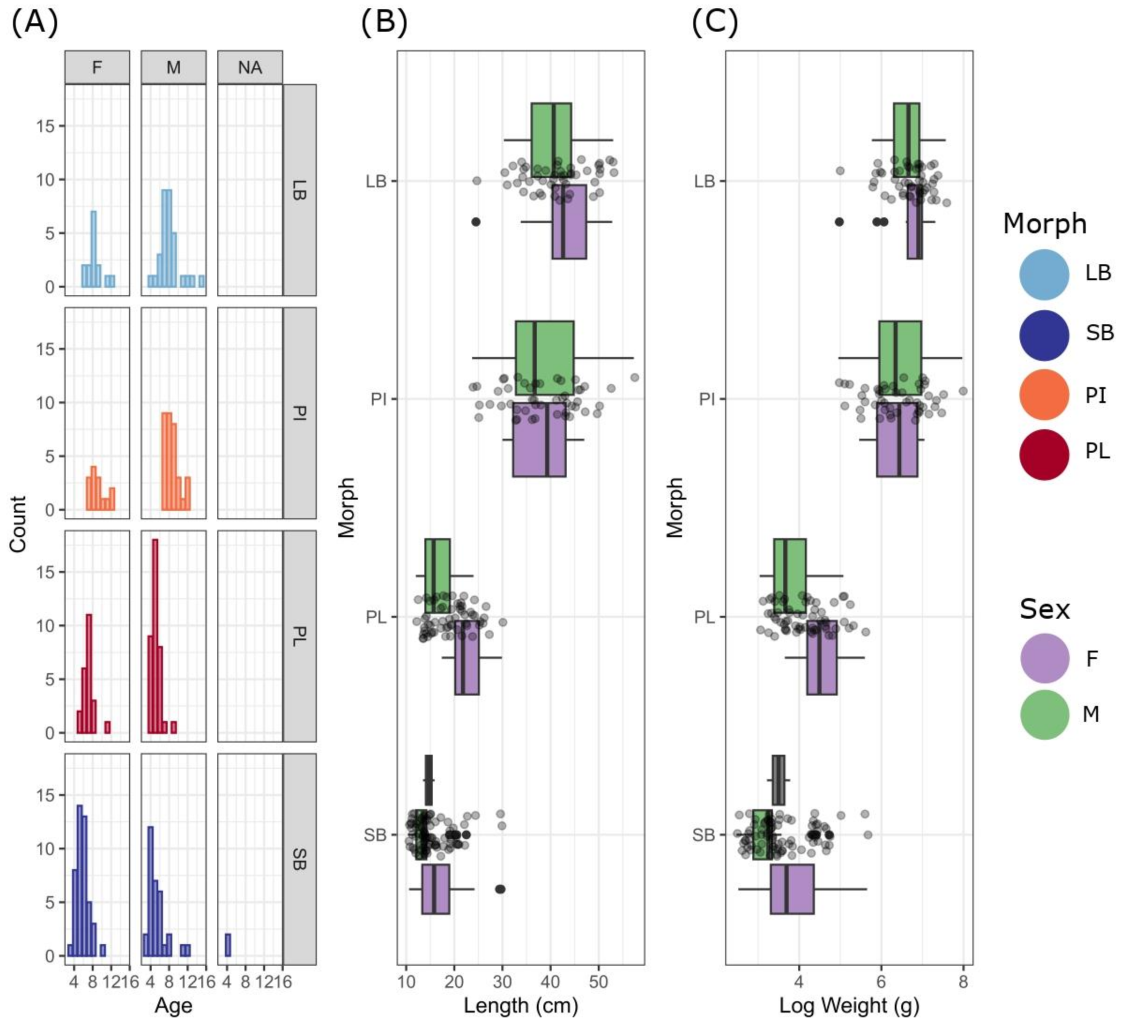


*S4 Appendix: Variation in size and age of sexually mature fish of the four sympatric morphs. (A) Histogram, showing age (years) distribution for all morphs by sex (NA indicates SB that could not be sexed. (B) The variation in fork length (FL, cm) by morph by sex represented by a boxplot. (C) Boxplot of the variation in* $log_e$ *weight (g) by morph by sex*.

*S5 Appendix: ANOVA results from tests of the influence of morph and sex effects on length (cm FL), $log_e$ weight (g) and age (years).*

| Dataset | | Morph | | Sex | | Morph x Sex | |
|---|---|---|---|---|---|---|---|
| | N | F | P | F | P | F | P |
| Fork length (cm) | 238 | 350.94 | << **0.001** | 14.39 | **0.0002** | 2.44 | 0.065 |
| $Log_e$ weight (g) | 238 | 381.28 | << **0.001** | 21.60 | < **0.001** | 3.95 | 0.009 |
| Age | 231 | 56.75 | << **0.001** | 8.41 | **0.004** | 3.00 | 0.032 |

F = F-statistic, P = P - value, < 0.05 in bold.

*S6 Appendix: Results from the Procrustes ANOVA, for effects of morph and $log_e$ FL on the size (centroid size) of the six bones.*

| Dataset/bone | $Log_e$ FL Effect | | | | Morph Effect | | | | Morph x $Log_e$ FL Interaction effect | | | |
|---|---|---|---|---|---|---|---|---|---|---|---|---|
| | $R^2$ | F | Z | P | $R^2$ | F | Z | P | $R^2$ | F | Z | P |
| Dentary | 0.959 | 8103.99 | 16.3 | **0.001** | 0.01 | 29.5 | 6.81 | **0.001** | 0.003 | 8.26 | 3.67 | **0.001*** |
| Articular-angular | 0.967 | 9171.03 | 16.62 | **0.001** | 0.005 | 16.36 | 5.7 | **0.001** | 0.003 | 9.88 | 3.98 | **0.001*** |
| Quadrat | 0.966 | 9580.49 | 16.36 | **0.001** | 0.008 | 27.60 | 7.23 | **0.001** | 0.003 | 8.37 | 4.04 | **0.001*** |
| Maxilla | 0.964 | 8203.91 | 16.98 | **0.001** | 0.006 | 16.8 | 5.11 | **0.001** | 0.003 | 8.35 | 3.65 | **0.001*** |
| Premaxilla | 0.936 | 6398.61 | 15.73 | **0.001** | 0.026 | 59.00 | 8.51 | **0.001** | 0.004 | 8.99 | 3.90 | **0.001*** |
| Supramaxilla | 0.96 | 6879.78 | 14.84 | **0.001** | 0.008 | 18.3 | 5.53 | **0.001** | 0.001 | 2.77 | 1.66 | 0.05 |

* Despite the p-values, all $R^2$ for the interaction term were small and no pairwise test of slope by morphs were significant.

$R^2$ = coefficient of determination; F = F-statistic; Z = Z-statistic; P = P-value, < 0.008 in bold.

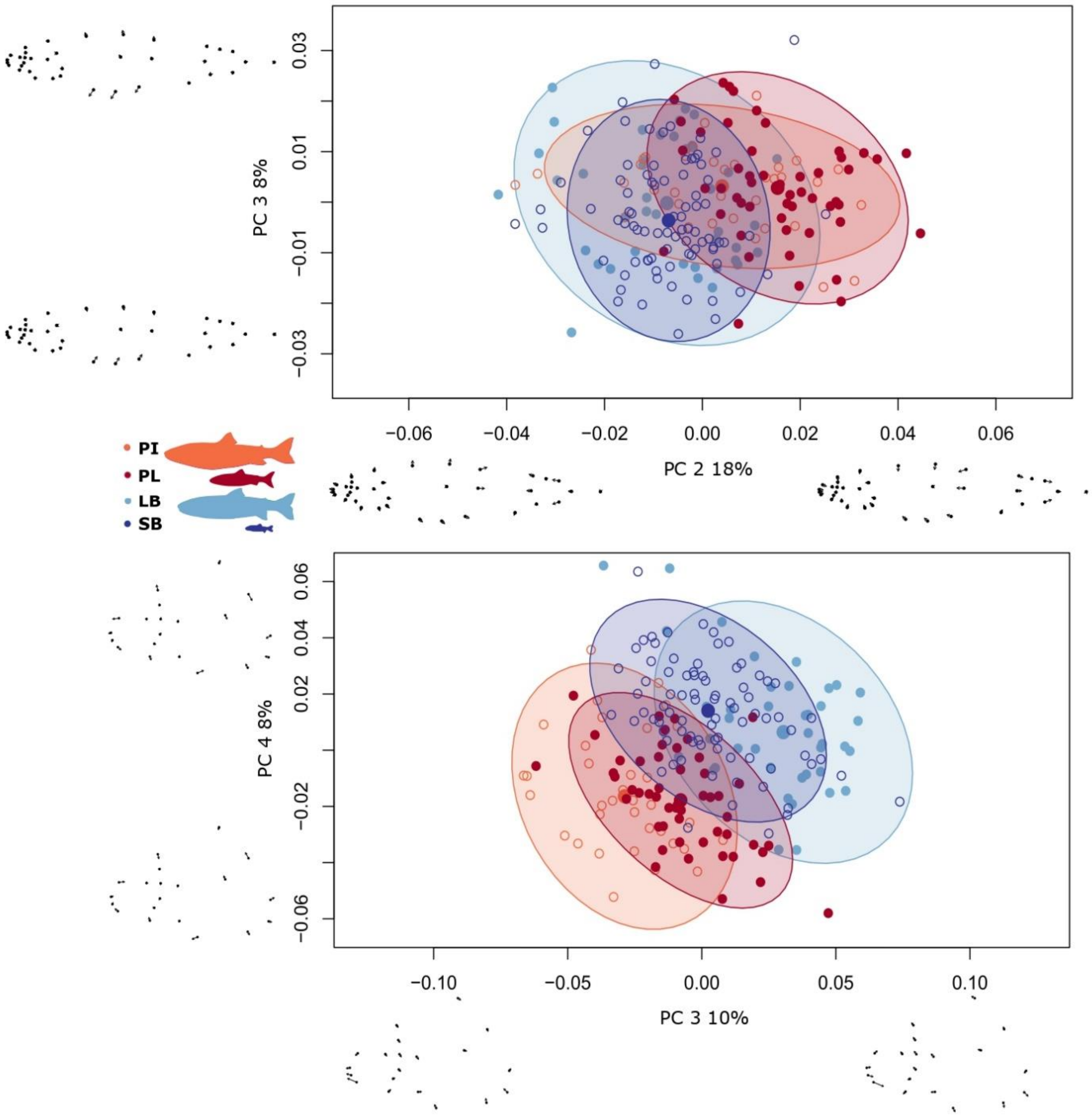


*S7 Appendix: Size corrected PC-plots of specimens and deformation grids showing shape variation in (top) external whole-body shape and (bottom) head shape. Plots of shape warps on X- and Y-axis are unmagnified. Each dot represents an individual and the ellipses represent 95% CI for the distribution by morph (large dot represents the mean of each morph distribution in these two dimensions of shape). For the whole-body (top) PC2 and 3 explain 18% and 8% of the variation respectively (PC1 was biased by sampling error, and not depicted) and for the head shape (bottom) PC3 and 4 explain 40% and 22% respectively (PC1 and PC2 were biased by sampling error, and not depicted).*

*S8 Appendix: Shape variation in the two external shapes and 6 head bones, analysed with Procrustes ANOVA, testing for influence of sex and size (centroid size).*

| Data | (a) $Log_e$ CS | | | | (b) Sex | | | |
|---|---|---|---|---|---|---|---|---|
| | $R^2$ | F | Z | P | $R^2$ | F | Z | P |
| Whole-body | 0.157 | 39.59 | 6.90 | **0.001** | 0.037 | 9.28 | 4.19 | **0.001** |
| Head | 0.206 | 53.34 | 7.02 | **0.001** | 0.012 | 3.03 | 2.48 | **0.007** |
| Dentary | 0.235 | 73.98 | 5.35 | **0.001** | 0.015 | 4.87 | 2.64 | **0.003** |
| Articular-angular | 0.189 | 55.71 | 5.88 | **0.001** | 0.011 | 3.36 | 2.56 | **0.004** |
| Quadrat | 0.300 | 102.22 | 6.27 | **0.001** | 0.009 | 3.14 | 2.42 | **0.008** |
| Premaxilla | 0.165 | 47.93 | 7.76 | **0.001** | 0.023 | 6.58 | 4.37 | **0.001** |
| Maxilla | 0.198 | 59.45 | 7.83 | **0.001** | 0.019 | 5.65 | 3.36 | **0.001** |
| Supramaxilla | 0.098 | 25.16 | 6.10 | **0.001** | 0.018 | 4.59 | 2.80 | **0.003** |

R2 = coefficient of determination; F = F-statistic; Z = Z-statistic; P = P-value, ≤ 0.008 in bold.

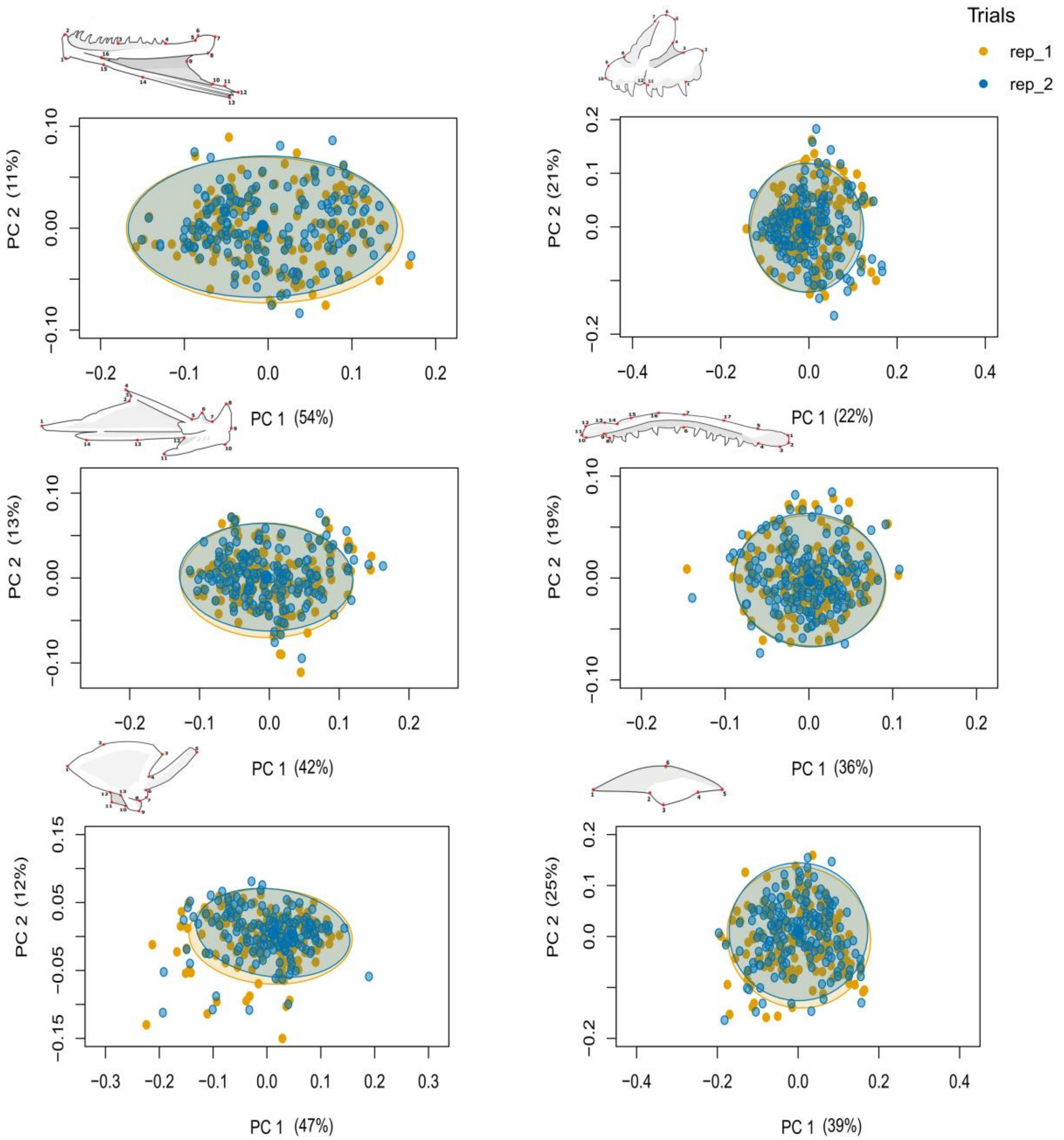


*S9 Appendix: PC-plots of specimens showing shape differences between replicate scoring on 168 specimens (rep1 and rep2), for dentary, premaxilla, articular-angular, maxilla, quadrat and supramaxilla (from top to bottom, left to right). Each dot represents an individual and the ellipses represent 95% CI for the distribution by replicates (large dot represents the mean for each morph replicates in these dimensions).*

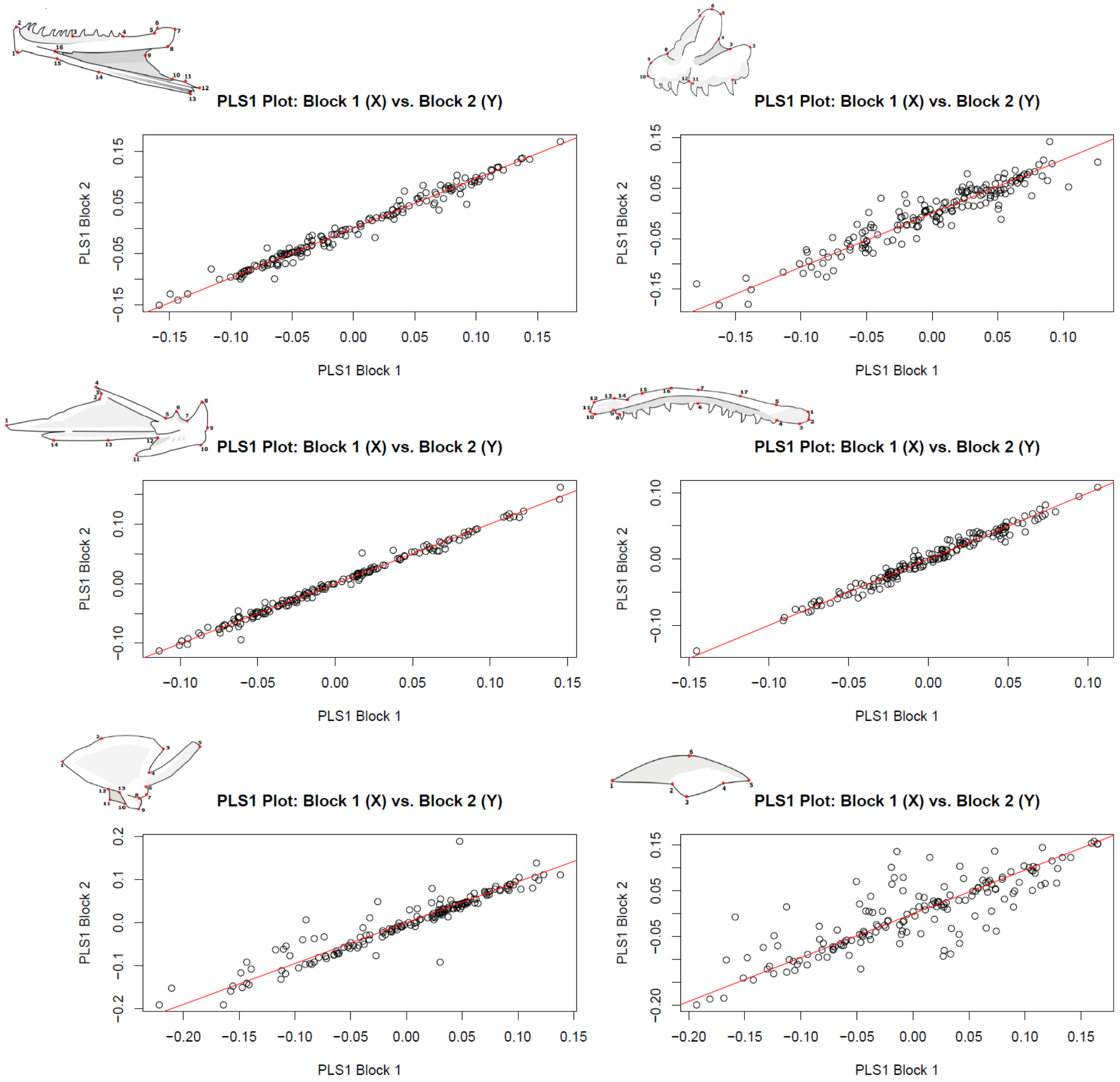


*S10 Appendix: Ordination plot (from partial least squares analysis (PLS)) showing the degree of association between the two replicates (trials) for dentary, premaxilla, articular-angular, maxilla, quadrat and supramaxilla (from top to bottom, left to right). Block 1 (x-axis) is replicate 1 and Block 2 (y-axis) is replicate 2. For all bones association between the replicates was always significant ($p < 0.001$). With the Pearson correlation coefficient being, dentary: 0.989, premaxilla: 0.939, articular-angular: 0.994, maxilla: 0.986, quadrat: 0.942 and supramaxilla: 0.835.*

*S11 Appendix: ANOVA results from tests of the influence of individual variation and measurement error (replicates) effects on shape variation in 6 jaw bones from 168 individuals, analysed with Procrustes ANCOVA.*

| Dataset/bone | Individual (id) | | | | Replicates (rep) | | | | Residuals |
|---|---|---|---|---|---|---|---|---|---|
| | $R^2$ | F | Z | P | $R^2$ | F | Z | P | $R^2$ |
| Dentary | 0.948 | 20.00 | 25.60 | **0.001** | 0.004 | 14.95 | 4.66 | **0.001** | 0.047 |
| Articular-angular | 0.935 | 14.85 | 23.85 | **0.001** | 0.002 | 4.63 | 3.04 | **0.002** | 0.063 |
| Quadrate | 0.929 | 14.01 | 23.65 | **0.001** | 0.004 | 11.16 | 4.92 | **0.001** | 0.066 |
| Maxilla | 0.935 | 15.57 | 21.60 | **0.001** | 0.005 | 13.29 | 4.81 | **0.001** | 0.060 |
| Premaxilla | 0.898 | 9.15 | 28.92 | **0.001** | 0.004 | 7.24 | 4.92 | **0.001** | 0.098 |
| Supramaxilla | 0.901 | 9.34 | 14.70 | **0.001** | 0.002 | 3.92 | 2.32 | 0.008 | 0.096 |

$R^2$ = coefficient of determination; F = F-statistic; Z = Z-statistic; P = P-value, < 0.008 in bold.

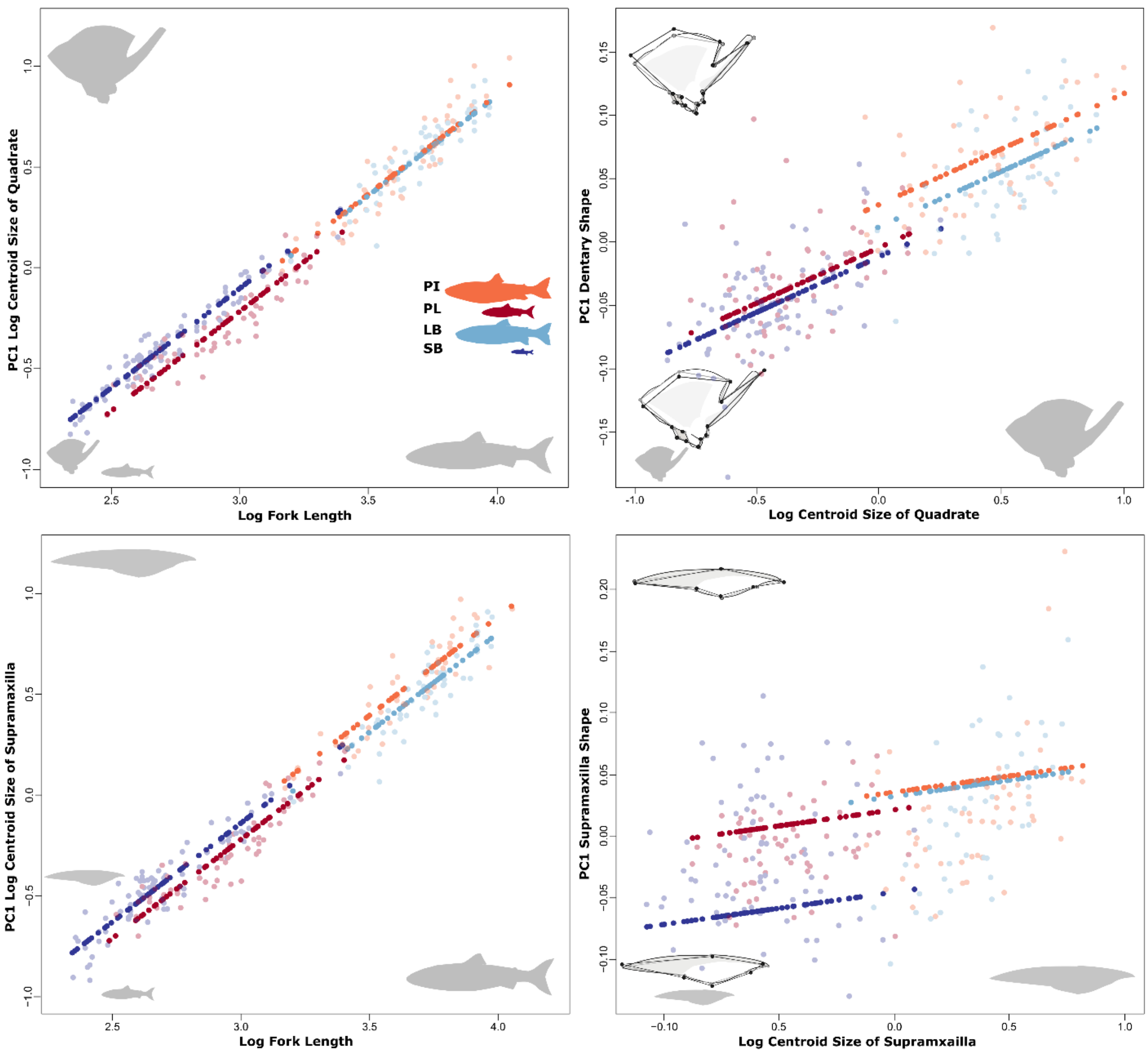


*S12 Appendix: Size and shape allometry of (top) quadrate and (bottom) supramaxilla). Left panels: relationships between bone size and fork length by morph; Right panels: relationships between bone shape and bone size. On the right are inset the associated shape changes related to each component, grey outlines the mean shape, and black the extremes for each PC. Shown are values for individuals (open circles) and the predicted values (filled) of regressions for both bone size ($CS_{bone}$) vs body size ($log_e$ FL) or bone-shape vs bone-size. $Log_e$, natural log transformation.*

*S13 Appendix: Estimates of regression for the relationships between the size of the individual and the size of the bone, per morph. See also Fig 2 and S6 and S12 Appendix.*

| Bones | Morphs | Intercept | Slope | $R^2$ |
|---|---|---|---|---|
| Dentary | LB | -2.86 | 1.19 | 0.97 |
| | SB | -1.9 | 0.91 | 0.97 |
| | PL | -1.77 | 0.87 | 0.97 |
| | PI | -2.59 | 1.16 | 0.97 |
| Articular-angular | LB | -3.02 | 1.17 | 0.97 |
| | SB | -2.09 | 0.92 | 0.97 |
| | PL | -1.83 | 0.81 | 0.97 |
| | PI | -2.67 | 1.1 | 0.97 |
| Quadrate | LB | -3.38 | 1.05 | 0.98 |
| | SB | -3.06 | 0.97 | 0.98 |
| | PL | -2.83 | 0.85 | 0.98 |
| | PI | -3.84 | 1.19 | 0.98 |
| Premaxilla | LB | -4.31 | 1.2 | 0.96 |
| | SB | -3.52 | 0.99 | 0.96 |
| | PL | -3.53 | 0.92 | 0.96 |
| | PI | -4.75 | 1.34 | 0.96 |
| Maxilla | LB | -2.72 | 1.17 | 0.97 |
| | SB | -1.96 | 0.95 | 0.97 |
| | PL | -1.66 | 0.83 | 0.97 |
| | PI | -2.59 | 1.16 | 0.97 |
| Supramaxilla | LB | -3.91 | 1.15 | 0.97 |
| | SB | -3.14 | 0.95 | 0.97 |
| | PL | -3.12 | 0.91 | 0.97 |
| | PI | -3.42 | 1.04 | 0.97 |

*S14 Appendix: Results from pairwise comparisons of variation in bone size between morphs for the six bones (dentary, articular-angular, quadrate, premaxilla, maxilla and supramaxilla), based on a model assuming no difference in allometry by morphs.*

| Bones | Morph pairs | D-value | UCL | Z | P-value |
|---|---|---|---|---|---|
| Dentary | LB-PI | 0.152 | 0.044 | 4.46 | **0.001** |
| | SB-PL | 0.003 | 0.036 | -1.13 | 0.854 |
| | LB-PL | 0.0004 | 0.064 | -2.23 | 0.988 |
| | PI-SB | 0.155 | 0.069 | 3.23 | **0.001** |
| | LB-SB | 0.003 | 0.072 | -1.45 | 0.909 |
| | PI-PL | 0.152 | 0.059 | 3.54 | **0.001** |
| Articular-angular | LB-PI | 0.082 | 0.035 | 3.41 | **0.001** |
| | SB-PL | 0.046 | 0.031 | 2.45 | **0.002** |
| | LB-PL | 0.027 | 0.049 | 0.59 | 0.3 |
| | PI-SB | 0.063 | 0.054 | 1.87 | 0.024 |
| | LB-SB | 0.019 | 0.059 | -0.07 | 0.543 |
| | PI-PL | 0.109 | 0.046 | 3.59 | **0.001** |
| Quadrate | LB-PI | 0.008 | 0.035 | -0.55 | 0.714 |
| | SB-PL | 0.115 | 0.032 | 4.55 | **0.001** |
| | LB-PL | 0.081 | 0.053 | 2.37 | 0.006 |
| | PI-SB | 0.026 | 0.057 | 0.38 | 0.370 |
| | LB-SB | 0.034 | 0.059 | 0.66 | 0.271 |
| | PI-PL | 0.088 | 0.05 | 2.83 | **0.001** |
| Premaxilla | LB-PI | 0.050 | 0.054 | 1.44 | 0.073 |
| | SB-PL | 0.216 | 0.045 | 5.58 | **0.001** |
| | LB-PL | 0.133 | 0.081 | 2.59 | **0.002** |
| | PI-SB | 0.033 | 0.085 | 0.18 | 0.436 |
| | LB-SB | 0.084 | 0.09 | 1.5 | 0.066 |
| | PI-PL | 0.183 | 0.075 | 3.56 | **0.001** |
| Maxilla | LB-PI | 0.08 | 0.039 | 3.17 | **0.001** |
| | SB-PL | 0.053 | 0.035 | 2.40 | 0.005 |
| | LB-PL | 0.056 | 0.060 | 1.5 | 0.064 |
| | PI-SB | 0.084 | 0.065 | 2.13 | 0.012 |
| | LB-SB | 0.003 | 0.07 | -1.43 | 0.913 |
| | PI-PL | 0.137 | 0.051 | 3.52 | **0.001** |
| Supramaxilla | LB-PI | 0.083 | 0.039 | 3.1 | **0.001** |
| | SB-PL | 0.081 | 0.036 | 3.22 | **0.001** |
| | LB-PL | 0.042 | 0.064 | 0.98 | 0.181 |
| | PI-SB | 0.044 | 0.068 | 0.86 | 0.198 |
| | LB-SB | 0.039 | 0.071 | 0.63 | 0.275 |

| | PI-PL | 0.125 | 0.057 | 3.2 | **0.002** |
|---|---|---|---|---|---|

D = pairwise distances between means; UCL = upper confidence level; Z = Z-statistic; P-value < 0.008 in bold (Bonferroni).

*S15 Appendix*: *Information on PCA, for all bone shapes, when size effects have not been removed. Symbols indicate how morphs align, # for when morphs align along morphotype, * when morphs align along the size- gradient. Double Symbols (* or #) for prominent separation and symbol in brackets for minor.*

| Bones | PCs | Eigenvalues | Proportion of Variance | Cumulative Proportion |
|---|---|---|---|---|
| Dentary | PC 1 ## | 0.0047 | 0.504 | 0.504 |
| | PC 2 * | 0.0012 | 0.128 | 0.632 |
| | PC 3 | 0.0006 | 0.062 | 0.694 |
| | PC 4 | 0.0006 | 0.059 | 0.752 |
| | PC 5 | 0.0004 | 0.045 | 0.797 |
| Articular-angular | PC 1 #* | 0.0033 | 0.404 | 0.404 |
| | PC 2 * | 0.0012 | 0.142 | 0.546 |
| | PC 3 | 0.0009 | 0.105 | 0.652 |
| | PC 4 | 0.0006 | 0.077 | 0.729 |
| | PC 5 | 0.0003 | 0.042 | 0.771 |
| Quadrate | PC 1 ** | 0.0047 | 0.456 | 0.457 |
| | PC 2 (*) | 0.0012 | 0.112 | 0.569 |
| | PC 3 | 0.0010 | 0.093 | 0.662 |
| | PC 4 | 0.0007 | 0.068 | 0.730 |
| | PC 5 | 0.0006 | 0.054 | 0.784 |
| Premaxilla | PC 1 ** | 0.0049 | 0.261 | 0.261 |
| | PC 2 ## | 0.0039 | 0.209 | 0.470 |
| | PC 3 | 0.0017 | 0.090 | 0.560 |
| | PC 4 | 0.0015 | 0.078 | 0.638 |
| | PC 5 | 0.0013 | 0.066 | 0.704 |
| Maxilla | PC 1 ** | 0.0018 | 0.345 | 0.345 |
| | PC 2 (*) | 0.0013 | 0.242 | 0.587 |
| | PC 3 # | 0.0007 | 0.131 | 0.718 |
| | PC 4 | 0.0003 | 0.054 | 0.771 |
| | PC 5 | 0.0002 | 0.045 | 0.816 |
| Supramaxilla | PC 1 | 0.0060 | 0.381 | 0.381 |
| | PC 2 * | 0.0040 | 0.252 | 0.633 |
| | PC 3 | 0.0030 | 0.189 | 0.823 |
| | PC 4 (*) | 0.0016 | 0.102 | 0.925 |
| | PC 5 | 0.0005 | 0.031 | 0.956 |

*S16 Appendix: Pairwise distances from tests of dentary shape allometry by morph, between vector angles and absolute differences between vector lengths, following ANCOVA model with the interaction of size (bone centroid size) and morph.*

| Test | Morph pairs | *r* | *angle* | UCL (95%) | Z | P-value |
|---|---|---|---|---|---|---|
| Vector angles | LB-PI | 0.420 | 65.153 | 44.882 | 3.09 | **0.003** |
| | SB-PL | 0.921 | 22.972 | 42.250 | -0.85 | 0.823 |
| | LB-PL | 0.781 | 38.657 | 47.151 | 0.86 | 0.197 |
| | PI-SB | 0.513 | 59.107 | 38.999 | 3.38 | **0.001** |
| | LB-SB | 0.626 | 51.210 | 42.509 | 2.40 | 0.010 |
| | PI-PL | 0.470 | 61.942 | 44.132 | 3.04 | **0.003** |
| | | | *d* | UCL (95%) | Z | P-value |
| Vector lengths | LB-PI | - | 0.071 | 0.050 | 2.28 | **0.003** |
| | SB-PL | - | 0.043 | 0.041 | 1.66 | 0.037 |
| | LB-PL | - | 0.047 | 0.048 | 1.49 | 0.061 |
| | PI-SB | - | 0.019 | 0.038 | 0.45 | 0.348 |
| | LB-SB | - | 0.090 | 0.045 | 3.02 | **0.001** |
| | PI-PL | - | 0.024 | 0.045 | 0.66 | 0.279 |

*r* = slope vector correlations; *angle:* between the two vectors being compared; *d* = pairwise distances between means; UCL = upper confidence level; Z = Z-statistic; P-value, ≤ 0.008 in bold (Bonferroni correction).

*S17 Appendix: Results from pairwise tests of shape by morphs, for differences between absolute vector lengths (D) used to examine possible shape differences by morph in five bones. These bones had significant shape differences according to ANOVA model with main effects and interaction of morph by bone size. Used to verify results from ANOVA and test for true allometry (i.e., different group slopes).*

| Bones | Morph pairs | D | UCL | Z | P-value |
|---|---|---|---|---|---|
| Dentary | LB-PI | 0.071 | 0.050 | 2.28 | **0.003** |
| | PL-SB | 0.043 | 0.041 | 1.66 | 0.037 |
| | LB-PL | 0.047 | 0.048 | 1.49 | 0.061 |
| | PI-SB | 0.019 | 0.038 | 0.45 | 0.348 |
| | LB-SB | 0.090 | 0.045 | 3.02 | **0.001** |
| | PI-PL | 0.024 | 0.045 | 0.66 | 0.279 |
| Articular-angular | LB-PI | 0.076 | 0.046 | 2.61 | **0.001** |
| | PL-SB | 0.021 | 0.044 | 0.53 | 0.322 |
| | LB-PL | 0.087 | 0.047 | 2.86 | **0.001** |
| | PI-SB | 0.032 | 0.039 | 1.23 | 0.116 |
| | LB-SB | 0.108 | 0.043 | 3.52 | **0.001** |
| | PI-PL | 0.010 | 0.045 | -0.35 | 0.635 |
| Premaxilla | LB-PI | 0.031 | 0.058 | 0.59 | 0.295 |
| | PL-SB | 0.004 | 0.053 | -1.23 | 0.877 |
| | LB-PL | 0.070 | 0.058 | 1.92 | 0.023 |
| | PI-SB | 0.044 | 0.048 | 1.40 | 0.070 |
| | LB-SB | 0.075 | 0.060 | 2.03 | 0.010 |
| | PI-PL | 0.039 | 0.054 | 1.06 | 0.157 |
| Maxilla | LB-PI | 0.035 | 0.038 | 1.42 | 0.083 |
| | PL-SB | 0.001 | 0.034 | -1.91 | 0.970 |
| | LB-PL | 0.049 | 0.039 | 2.08 | 0.010 |
| | PI-SB | 0.015 | 0.030 | 0.54 | 0.333 |
| | LB-SB | 0.050 | 0.037 | 2.16 | 0.011 |
| | PI-PL | 0.014 | 0.036 | 0.28 | 0.410 |
| Supramaxilla | LB-PI | 0.079 | 0.078 | 1.60 | 0.049 |
| | PL-SB | 0.010 | 0.068 | -0.74 | 0.771 |
| | LB-PL | 0.116 | 0.077 | 2.43 | 0.002 |
| | PI-SB | 0.047 | 0.067 | 1.00 | 0.171 |
| | LB-SB | 0.126 | 0.081 | 2.49 | **0.004** |
| | PI-PL | 0.037 | 0.073 | 0.59 | 0.290 |

D = pairwise differences between vector lengths; UCL = upper confidence level; Z = Z-statistic; P-value, < 0.008 in bold.

*S18 Appendix: Results from "Significant Vector Angle test" examining possibly allometric shape differences by morph for five bones, that showed significant allometric differences according to ANOVA model with main and interaction effects (morph x bone size). Used to verify results from ANOVA and test for true allometry (i.e., different group slopes).*

| Bones | Morph pairs | R | Angle | UCL | Z | P-value |
|---|---|---|---|---|---|---|
| Dentary | LB-PI | 0.420 | 65.153 | 44.882 | 3.09 | **0.003** |
| | PL-SB | 0.921 | 22.972 | 42.250 | -0.85 | 0.823 |
| | LB-PL | 0.781 | 38.657 | 47.151 | 0.86 | 0.197 |
| | PI-SB | 0.513 | 59.107 | 38.999 | 3.38 | **0.001** |
| | LB-SB | 0.626 | 51.210 | 42.509 | 2.40 | 0.010 |
| | PI-PL | 0.470 | 61.942 | 44.132 | 3.04 | **0.003** |
| Articular-angular | LB-PI | 0.840 | 32.810 | 46.375 | 0.17 | 0.431 |
| | PL-SB | 0.857 | 31.044 | 44.361 | 0.18 | 0.420 |
| | LB-PL | 0.662 | 48.522 | 48.129 | 1.63 | 0.046 |
| | PI-SB | 0.770 | 39.629 | 41.350 | 1.50 | 0.067 |
| | LB-SB | 0.744 | 41.952 | 44.675 | 1.39 | 0.085 |
| | PI-PL | 0.662 | 48.550 | 45.035 | 2.01 | 0.021 |
| Premaxilla | LB-PI | 0.623 | 51.493 | 60.860 | 0.86 | 0.199 |
| | PL-SB | 0.654 | 49.123 | 56.735 | 0.93 | 0.185 |
| | LB-PL | 0.531 | 57.960 | 65.830 | 1.07 | 0.158 |
| | PI-SB | 0.114 | 83.440 | 52.954 | 4.18 | **0.001** |
| | LB-SB | 0.079 | 85.480 | 62.032 | 3.00 | **0.001** |
| | PI-PL | 0.475 | 61.594 | 58.634 | 1.88 | 0.029 |
| Maxilla | LB-PI | 0.545 | 56.948 | 71.236 | 0.90 | 0.189 |
| | PL-SB | 0.611 | 52.346 | 64.807 | 0.97 | 0.178 |
| | LB-PL | 0.728 | 43.227 | 73.504 | -0.09 | 0.525 |
| | PI-SB | 0.124 | 82.870 | 59.338 | 2.78 | **0.003** |
| | LB-SB | 0.453 | 63.051 | 66.483 | 1.50 | 0.066 |
| | PI-PL | 0.546 | 56.873 | 66.917 | 1.07 | 0.153 |
| Supramaxilla | LB-PI | 0.951 | 17.948 | 102.730 | -2.39 | 0.992 |
| | PL-SB | -0.350 | 110.507 | 91.714 | 2.67 | **0.007** |
| | LB-PL | 0.217 | 77.454 | 99.746 | 0.80 | 0.234 |
| | PI-SB | -0.165 | 99.489 | 90.038 | 1.97 | 0.022 |
| | LB-SB | -0.188 | 100.864 | 95.853 | 1.83 | 0.025 |
| | PI-PL | 0.420 | 65.196 | 99.472 | 0.41 | 0.346 |

R = slope of vector correlation; UCL = upper confidence level; Z = Z-statistic; P-value < 0.008 in bold.

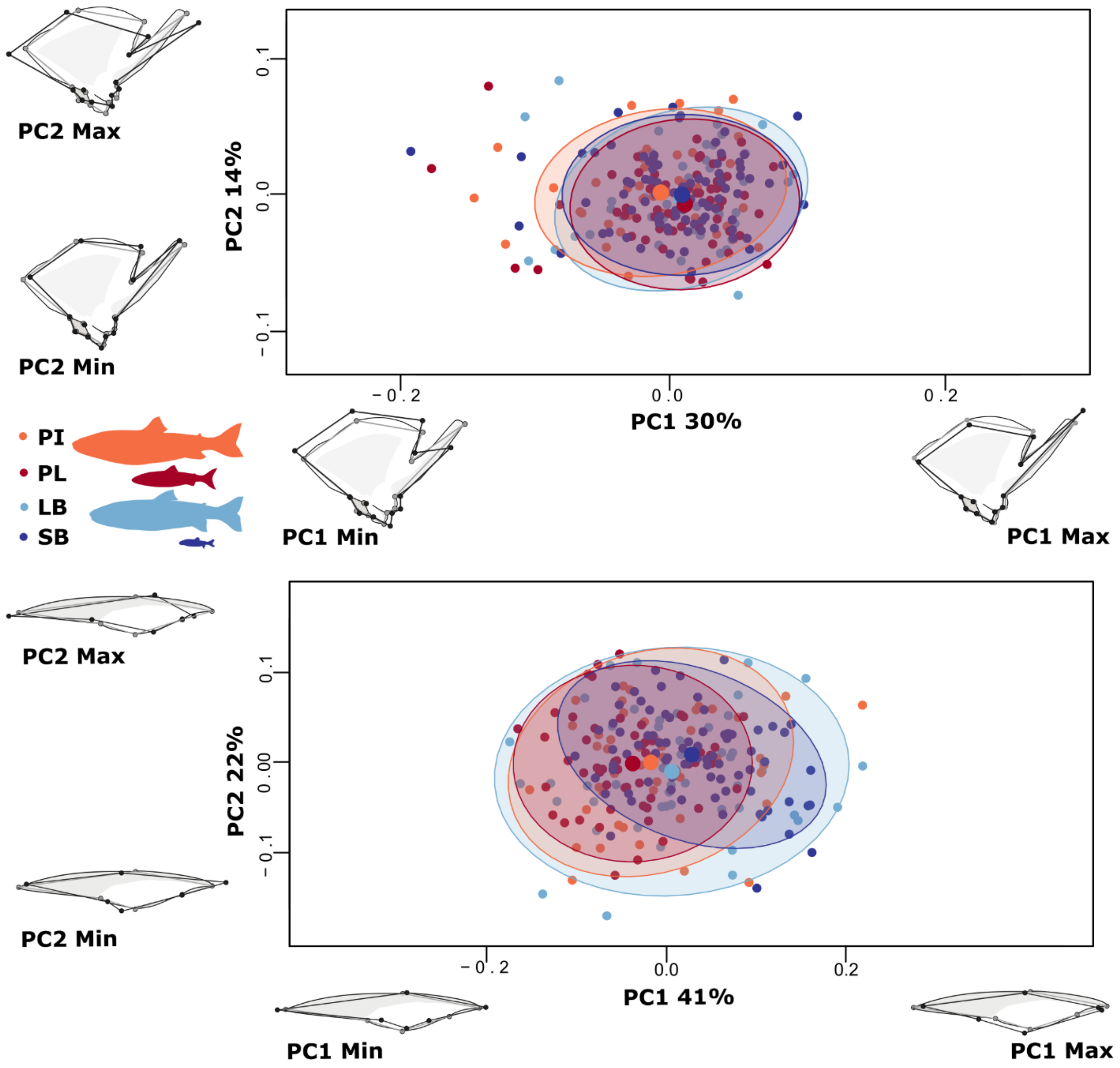


*S19 Appendix: Size corrected PC-plots of specimens and deformation grids showing shape variation in (top) quadrat and (bottom) supramaxilla. On each axis are the associated shape changes related to each component, grey outlines the mean shape and black the extremes for each PC. Plots of shape warps on X- and Y-axis are unmagnified. Each dot represents an individual and the ellipses represent 95% CI for the distribution by morph (large dot represents the mean of each morph distribution in these two dimensions of shape). For the quadrat (top) PC1 and 2 explain 30% and 14% of the variation respectively and the supramaxilla (bottom) PC1 and 2 explain 40% and 22% respectively.*

*S20 Appendix: Tests of integration among craniofacial bones in arctic charr. The six bones were compared using partial least squares (PLS), within anatomical regions (upper and lower jaws) and between regions.*

| Comparison | Bones | | r-PLS | Z | p-value |
|---|---|---|---|---|---|
| Lower Jaw | Dentary | Articular-angular | 0.917 | 7.081 | 0.001 |
| | Dentary | Quadrate | 0.663 | 5.244 | 0.001 |
| | Articular-angular | Quadrate | 0.692 | 6.243 | 0.001 |
| Upper Jaw | Premaxilla | Maxilla | 0.683 | 7.774 | 0.001 |
| | Premaxilla | Supramaxilla | 0.541 | 6.579 | 0.001 |
| | Maxilla | Supramaxilla | 0.682 | 11.384* | 0.001 |
| Across Jaws | Dentary | Premaxilla | 0.722 | 8.330 | 0.001 |
| | Articular-angular | Premaxilla | 0.724 | 7.388 | 0.001 |
| | Quadrate | Premaxilla | 0.694 | 6.063 | 0.001 |
| | Dentary | Maxilla | 0.775 | 6.917** | 0.001 |
| | Articular-angular | Maxilla | 0.745 | 6.845** | 0.001 |
| | Quadrate | Maxilla | 0.719 | 6.875 | 0.001 |
| | Dentary | Supramaxilla | 0.540 | 5.794 | 0.001 |
| | Articular-angular | Supramaxilla | 0.547 | 6.363 | 0.001 |
| | Quadrate | Supramaxilla | 0.550 | 5.410 | 0.001 |

*Effect size of maxilla-supramaxilla were significantly higher than premaxilla-maxilla.

**Effect size of dentary-maxilla and articular-angular-maxilla were significantly higher than articular-angular-supramaxilla.
